# Inhibition of Acid Sphingomyelinase Links Sphingolipid Remodeling to Necroptotic Cell Death

**DOI:** 10.64898/2026.03.10.710852

**Authors:** Luis Pilapil, Shweta Chitkara, G. Ekin Atilla-Gokcumen

## Abstract

Necroptosis is a lytic form of programmed cell death that requires activation of the RIPK1/3– MLKL complex and results in plasma membrane permeabilization. Although the protein components governing necroptosis are well defined, the lipid determinants of this process remain poorly understood. Here, we combined lipidomics, pharmacological perturbations of sphingolipid metabolism and functional assays to identify sphingolipid pathways that contribute to necroptotic cell death. Using a panel of small molecule inhibitors, we found that inhibition of acid sphingomyelinase (ASMase) with ARC39 restored cell viability and membrane integrity during necroptosis without altering canonical necroptotic signaling. Lipidomic analysis revealed that ARC39 treatment prevented ceramide accumulation in necroptosis, linking reduced ceramide levels to decreased membrane permeability. Interestingly, ARC39 treatment did not reduce total cellular levels of phosphorylated MLKL (pMLKL) nor its initial membrane association, suggesting that the observed decrease in membrane permeability arises downstream of MLKL activation. Instead, our findings support a model in which the reduction of ceramide levels impairs productive membrane insertion and pore formation by pMLKL. Consistent with this interpretation, genetic knockdown of ASMase similarly resulted in increased cell viability, decreased membrane permeabilization, and decreased ceramide levels during necroptosis, further linking ceramide homeostasis to necroptotic membrane damage. Together, these results indicate that ASMase-derived ceramides are important for efficient MLKL-mediated membrane permeabilization in necroptosis.

## Introduction

Necroptosis is a regulated form of programmed cell death that is distinguished by plasma membrane permeabilization and the subsequent release of intracellular contents, driving inflammatory responses.^1^ Unlike apoptosis, which preserves membrane integrity, necroptosis results in lytic membrane rupture as its execution step. Mechanistically, necroptotic signaling leads to the activation of the Receptor-interacting serine/threonine-protein kinase1/3 (RIPK1/3) and formation of the necrosome complex. RIPK3 consequently phosphorylates the pseudokinase, mixed lineage kinase-like (MLKL), triggering MLKL oligomerization and conformational activation.^1–3^ Activated MLKL translocates to the plasma membrane, where it associates with phospholipids, undergoes further assembly, and disrupts membrane integrity, ultimately resulting in permeabilization and cell lysis.^4^ Although the core RIPK1/3-MLKL signaling axis has been extensively characterized, the membrane and lipid determinants that enable MLKL to execute plasma membrane rupture remain incompletely understood.

Lipids can participate actively in necroptotic regulation by altering membrane biophysical characteristics, controlling membrane organization, and modulating protein-membrane interactions.^5^ Several lipid classes, including phosphatidylinositol phosphates, phosphatidylserines, and sphingolipids, are compositionally regulated during necroptosis, and these changes have been linked to signaling, membrane dynamics, and pore formation.^5, 6^

Ceramides have increasingly been implicated in necroptosis, both as bioactive signaling molecules and as structural regulators of membrane organization. Early evidence demonstrated that exogenous delivery of short-chain ceramides can directly induce necroptotic phenotypes. For example, treatment of head and neck squamous carcinoma cells with short chain C2-ceramides triggered programmed necrosis.^7^ Similarly, ovarian luteal tissue has been reported to undergo necroptotic cell death driven by the accumulation of endogenous ceramides generated through the salvage pathway.^8^ In this system, salvage pathway-derived ceramides promote necroptosis, as supported by the ability of exogenous ceramide to induce cell death and by the observed sensitivity of cell death to both the pan-caspase inhibitor and the MLKL oligomerization inhibitor.^8^

Consistent with these observations, a previous study showed that ceramides accumulated during necroptosis. More specifically, in an analysis comparing the sphingolipid composition of necroptotic to non-necroptotic cells, C16:0 ceramide was found to be selectively enriched in necroptotic cells.^9^ This finding was strengthened through the use of necroptosis-resistant sublines, enabling discrimination between lipid changes unique to necroptosis versus those shared with apoptosis.^9^

To define global lipid remodeling during necroptosis, we previously performed untargeted lipidomics in HT-29 cells and observed broad accumulation of multiple ceramide species.^10^ Targeted sphingolipidomics confirmed coordinated remodeling across ceramides, dihydroceramides, sphingomyelins, and glycosphingolipids.^10^ Mechanistically, we found that transcripts encoding ceramide synthases were elevated, implicating enhanced *de novo* biosynthesis as one contributor to increased ceramide levels during necroptosis.^10^ Together, these studies suggest that ceramide elevation can contribute to necroptotic death programs.

Beyond accumulation, ceramides have been proposed to functionally interfere with necroptotic signaling machinery. For example, the delivery of ceramide nanoliposomes to ovarian cancer cells induced necroptosis, and knockdown of MLKL significantly attenuated cell death, suggesting a functional interaction between ceramides and MLKL during execution.^11, 12^ Interestingly, studies using the sphingosine-1-phosphate receptor (S1PR) antagonist, FTY720, have demonstrated that ceramide binding to inhibitor 2 of protein phosphatase 2A (I2PP2A) activates PP2A, promoting RIPK1 kinase activity and necrotic cell death.^13–16^ Subsequent work further revealed that FTY720 induces the formation of a ceramide-protein complex that assembles into ceramide-enriched membrane pores termed *ceramidosomes*.^17^ However, this pathway proceeds independently of canonical RIPK3/MLKL-driven necroptosis and does not follow canonical necroptotic signaling.^13^

Collectively, these findings establish that ceramides accumulate during necroptosis and can potentiate necroptotic signaling or membrane damage. However, whether ceramide remodeling is mechanistically required for canonical RIPK3/MLKL-dependent necroptotic execution, and how ceramides influence MLKL membrane engagement, remain unresolved. Given their ability to reorganize membrane architecture, generate ordered lipid domains, and modulate protein insertion, we hypothesized that sphingolipids, and ceramides in particular, can function as central biophysical regulators of necroptotic membrane permeabilization.

To define how sphingolipids and their metabolic enzymes functionally contribute to necroptosis, we used small molecule inhibitors targeting different enzymes in the sphingolipid metabolic pathways and investigated the consequences of the inhibition of these enzymes in necroptosis. Among these compounds, inhibition of acid sphingomyelinase (ASMase) with ARC39 produced a striking restoration of viability and plasma membrane integrity during necroptosis. ARC39 did not reduce MLKL phosphorylation, suggesting that the compound acts downstream of canonical necroptotic signaling and MLKL activation.

To determine the sphingolipid species responsible for ARC39-mediated necroptotic rescue, we conducted lipidomic profiling of necroptotic cells pre-treated with ARC39. We observed a robust reduction in ceramide accumulation, strongly implicating ASMase-dependent ceramide generation in necroptotic membrane permeabilization. These findings are consistent with the idea that ceramide-rich membrane domains may facilitate the terminal stages of necroptosis and that their disruption mitigates MLKL-mediated membrane damage.^17, 18^ In this context, ARC39 likely disrupts a critical ASMase-derived ceramide pool necessary for membrane permeabilization. Together, our results identify sphingomyelinase-mediated ceramide production, particularly the ASMase-dependent pathway, as a key lipid regulatory mechanism governing necroptotic execution.

## Results and Discussion

### Inhibition of ASMase restores membrane permeability in necroptosis

Prior studies established that ceramides accumulate during necroptosis^9, 10^, prompting us to investigate whether specific steps within sphingolipid metabolism functionally regulate this process. To systematically interrogate the contribution of sphingolipid metabolic pathways to necroptotic execution, we assembled a panel of 13 inhibitors that targets key enzymes in the *de novo* ceramide biosynthesis, sphingomyelin hydrolysis, and downstream sphingolipid conversion processes (**Figure 1; Table S1**).

**Figure 1.**
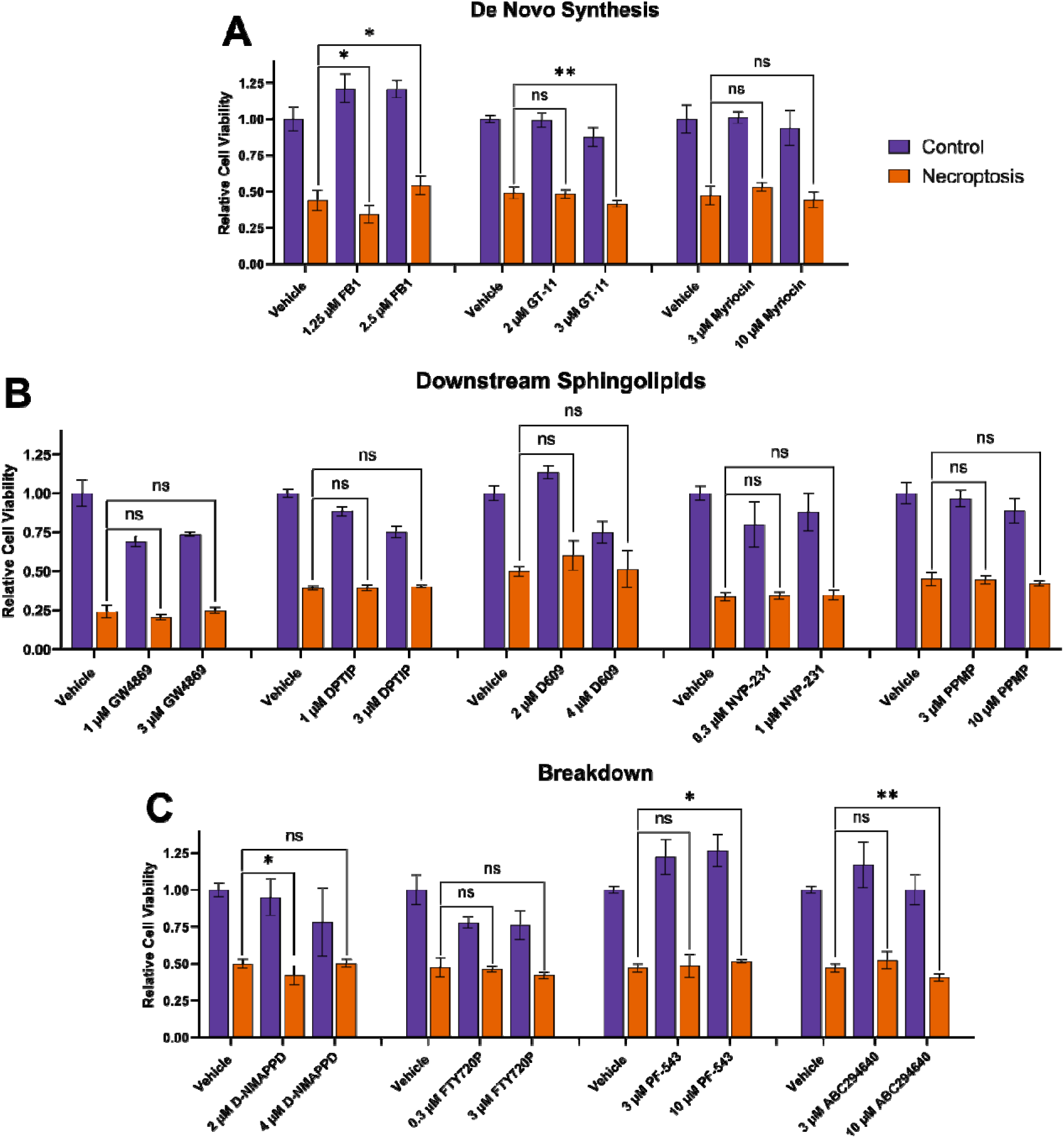
Cell viability measurements to study the involvement of sphingolipid pathway-related proteins in necroptotic signaling. HT-29 cells were pre-treated with inhibitors at the indicated concentrations for 21 hours, followed by the induction of necroptosis for 3 hours. Cell viability was subsequently assessed using the MTT assay. **(A)** Inhibition of the *de novo* ceramide biosynthetic pathway by FB1, GT-11, and Myriocin did not result in substantial changes in cell viability under necroptotic conditions. **(B)** Inhibitors targeting downstream sphingolipids (GW4869, DPTIP, D609, NVP-231, and PPMP) showed no significant change in cell viability compared to the vehicle control under necroptotic conditions. **(C)** Inhibitors targeting the sphingolipid breakdown pathways (D-NMAPPD, FTY720P, PF-543, and ABC29640) did not result in profound changes in cell viability relative to the vehicle control. Data represents the mean ± standard deviation; *n* = 5, ns represents *p* > 0.05, * represents *p* < 0.05, ** represents *p* < 0.01, *** represents *p* < 0.001.

We evaluated the impact of these perturbations on necroptotic cell death in HT29 human colorectal carcinoma cells. Necroptosis was induced using the established combination of zVAD-FMK, BV6, and TNF-α, and cell viability was assessed using the MTT assay which measures mitochondrial activity.^19, 20^ As expected, necroptosis induction produced a pronounced reduction in cell viability relative to untreated controls. To investigate the role of sphingolipid-related enzymes in this process, cells were pre-treated with inhibitors followed by necroptosis induction, and cell viability was assessed following necroptotic stimulation. Compounds were used at nanomolar to low micromolar concentrations (300 nM–10 µM) to minimize nonspecific cytotoxicity that could confound interpretation through off-target death pathways.

Given prior evidence that ceramides accumulate during necroptosis,^9, 10^ we first tested whether inhibition of *de novo* ceramide biosynthesis would exert a protective effect on necroptotic cell death (**Figure 1A**). We targeted key enzymes within the *de novo* ceramide biosynthetic pathway, including serine palmitoyltransferase using myriocin,^21, 22^ ceramide synthase using FB1,^23, 24^ and dihydroceramide desaturase using GT-11.^25^ Inhibition of the most upstream enzyme, serine palmitoyltransferase, did not rescue necroptotic cell death. Similarly, blocking ceramide synthase activity with FB1 produced only a modest increase in cell viability, observable primarily at higher concentrations. In contrast, inhibition of dihydroceramide desaturase with GT-11 slightly decreased viability under necroptotic conditions. Collectively, these findings indicate that *de novo* ceramide biosynthesis is not a major contributor to necroptosis.

We next examined whether metabolic pathways that convert ceramides into complex sphingolipids influence necroptotic execution (**Figure 1B**). To target downstream sphingolipid remodeling, we utilized GW4869^26, 27^ and DPTIP^28, 29^ (neutral sphingomyelinase 2 inhibitor), D609 (sphingomyelin synthase inhibitor),^30, 31^ NVP-231 (ceramide kinase inhibitor),^32^ and PPMP (glucosylceramide synthase inhibitor).^33^ Inhibition of these enzymes did not produce significant changes in necroptotic cell viability (*p* > 0.05) at the concentrations tested.

To further evaluate ceramide turnover and signaling, we perturbed sphingolipid catabolic pathways using D-NMAPPD (ceramidase inhibitor),^34^ FTY720P (sphingosine-1-phosphate receptor antagonist),^35–37^ PF-543 (sphingosine kinase-1 inhibitor),^38, 39^ and ABC294640 (sphingosine kinase-2 inhibitor)^40–42^ (**Figure 1C**). The inhibition of sphingosine kinase signaling or sphingosine-1-phosphate receptor pathways produced minimal functional effects. Although PF-543 treatment resulted in a modest increase in viability (*p* < 0.05) and ABC294640 produced a small decrease (*p* < 0.01), the magnitude of these changes was limited, suggesting that these enzymes do not play a major role in necroptosis.

In contrast to the minimal phenotypes observed across biosynthetic, downstream remodeling, trafficking, and catabolic pathways, inhibition of ASMase with ARC39^43, 44^ (**Figure 2A**) produced a striking rescue in cell viability during necroptosis. We first identified this phenotype using the MTT assay (**Figure 2B**).^20, 45^ To validate our phenotypic observation using an additional viability readout, we performed an ATP-based luminescence assay (**Figure 2C**),^46^ which likewise demonstrated a robust restoration of cellular viability upon ARC39 treatment.

**Figure 2.**
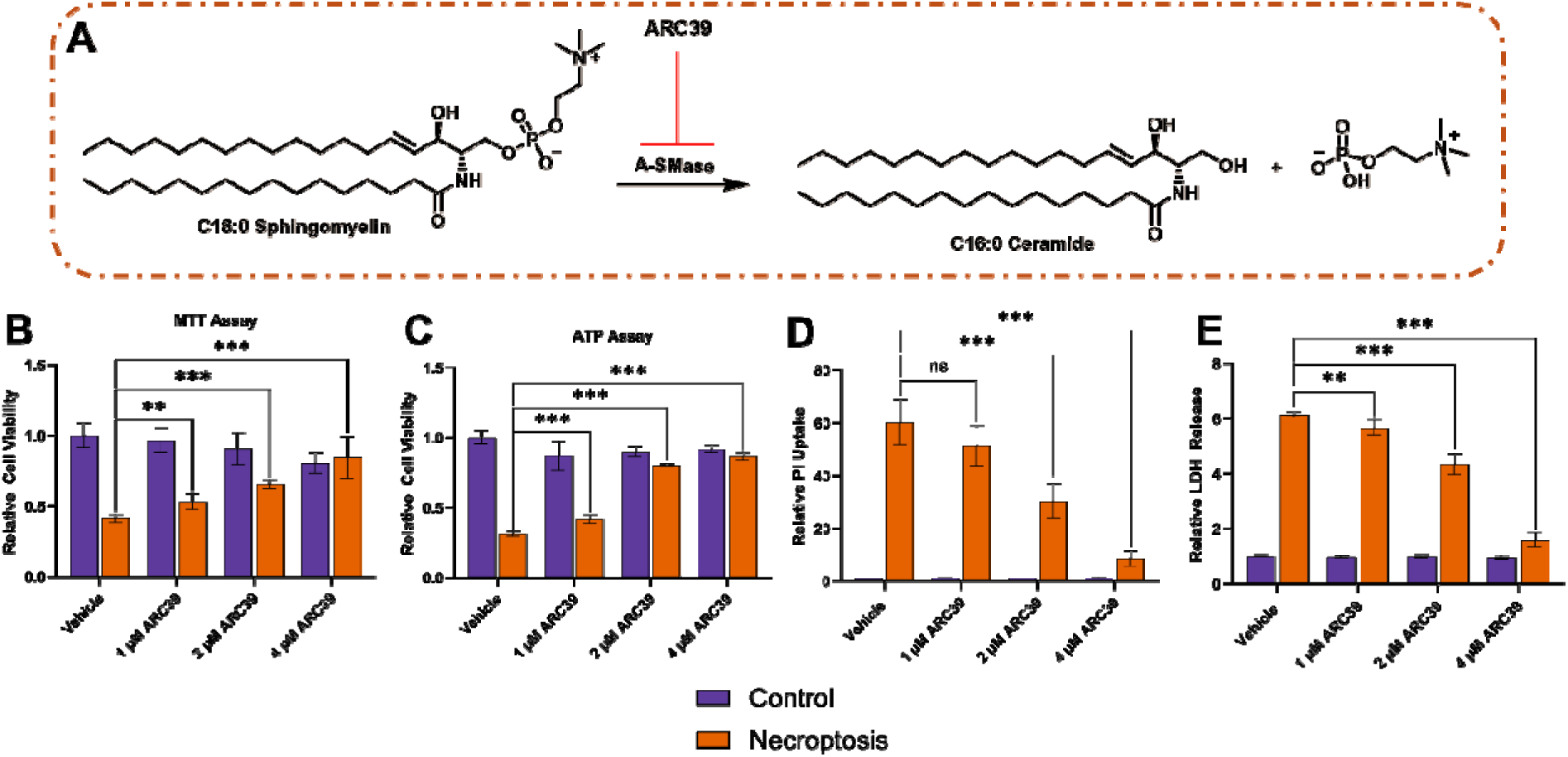
Treatment of ARC39 in necroptotic HT-29 cells results in recovery of of cell death and membrane permeability. **(A)** ARC39 inhibits ASMase, which catalyzes the hydrolysis of sphingomyelin into ceramide and phosphorylcholine. C16:0 sphingomyelin and C16:0 ceramide are shown as representative lipids for this reaction. **(B)** MTT assay data demonstrate that ARC39 treatment increases cell viability in a concentration-dependent manner under necroptotic conditions. **(C)** ATP assay results further confirm increased cell viability in a concentration-dependent manner compared to the vehicle control under necroptotic conditions. **(D)** PI assay data shows decreased PI uptake in ARC39-treated cells, indicating reduced membrane permeability. **(E)** LDH release assay similarly show decreased relative LDH release, further confirming reduced membrane permeability. Data represents the mean ± standard deviation; *n* = 5, ns represents *p* > 0.05, ** represents *p* < 0.01, *** represents *p* < 0.001.

Given the magnitude of this protective effect, we next asked whether ASMase inhibition have an impact on necroptotic membrane integrity, a terminal downstream consequence of MLKL activation. We assessed plasma membrane permeabilization using two independent assays: lactate dehydrogenase (LDH) release and propidium iodide (PI) uptake.^20, 47^ Consistent with the cell viability measurements, ARC39 treatment markedly suppressed membrane damage, as evidenced by decreased PI uptake (**Figure 2D**) and reduced LDH release (**Figure 2E**). The concordance across viability and membrane permeability assays demonstrates that ASMase inhibition not only preserves cellular metabolic activity but also functionally protects plasma membrane integrity during necroptotic execution.

These findings collectively reveal an important observation regarding the role of the sphingolipid pathway in necroptosis. While perturbation of *de novo* biosynthesis, most downstream sphingolipid conversion, and catabolic signaling pathways showed minimal effect on necroptosis, inhibition of ASMase robustly attenuated necroptotic cell death. Our results suggest that ASMase-dependent ceramide generation is a critical regulator of necroptotic membrane permeabilization and that sphingomyelin hydrolysis is a previously unrecognized potential lipid checkpoint governing necroptotic execution.

### ASMase inhibition suppresses ceramide accumulation in necroptosis

Sphingomyelinases catalyze the hydrolysis of sphingomyelin to ceramide, and two major isoforms (ASMase and neutral sphingomyelinase 2 (NSMase2)) are primarily differentiated based upon their subcellular location and cation dependency.^48^ NSMase2 (encoded by *SMPD3*) is the predominant form of neutral sphingomyelinase that is involved in the cellular stress response and is dependent on Mg^2+^ and neutral pH for enzyme activity.^27, 49, 50^ It is also primarily associated with the plasma membrane^51^ and Golgi,^49^ where it can modulate local ceramide pools.^50, 52, 53^

On the other hand, ASMase is encoded by the *SMPD1* gene and can give rise two distinct enzymes, namely secreted ASMase or lysosomal ASMase.^48, 54–56^ Both forms originate from a common precursor protein but differ in their post-translational processing and trafficking. Secreted ASMase is released into the extracellular space via the secretory pathway, whereas lysosomal ASMase is targeted to the endolysosomal compartment. Importantly, lysosomal ASMase can translocate to the plasma membrane under stress conditions, where it hydrolyzes sphingomyelin to generate ceramide and promote membrane reorganization and signaling.

To probe the involvement of ASMase and ceramide production in necroptosis more directly, we performed targeted lipidomics on necroptotic cells treated with ARC39 and made comparisons with control and necroptotic cells (**Figure 3**). The lipid composition was analyzed by LC-MS using our established protocols.^57, 58^ We examined triacylglycerols, phosphatidylcholines and various sphingolipids which includes ceramides, dihydroceramides, sphingomyelins, and hexosylceramides.

**Figure 3.**
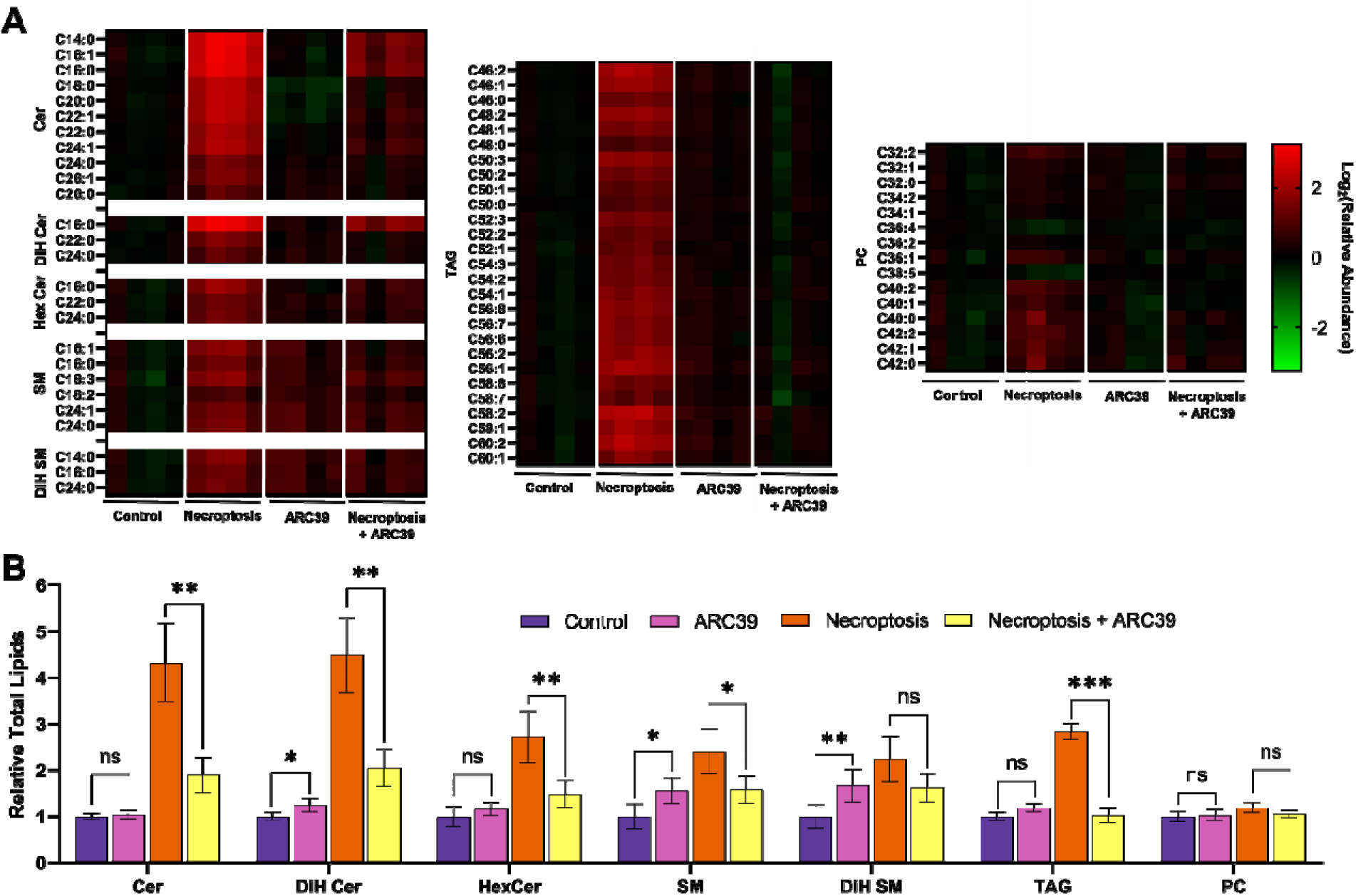
LC-MS analysis shows that ASMase inhibition significantly reduces ceramide levels during necroptosis. **(A)** Heatmap displaying the relative total lipid changes under control, 4 μM ARC39 treatment, necroptosis, and necroptosis with 4 μM ARC39 treatment. The relative abundance is presented as Log_2_(relative abundance). The relative abundance was calculated by normalizing each lipid species to the raw abundance of the internal standard. The internal standard corrected abundances were then normalized to the internal standard-corrected abundance of the respective control. Four biological replicates were analyzed per condition (n = 4). Red indicates accumulation, black indicates baseline levels, and green indicates depletion. Cer, Ceramide; DiHCer, Dihydroceramide; HexCer, Hexosylceramide; SM, Sphingomyelin; DiHSM, Dihydrosphingomyelin; TAG, Triacylglycerol; PC, Phosphatidylcholine. Internal standards used for correction: C17:0 Ceramide for Cer, DiH Cer and Hex Cer; C17:0 Sphingomyelin for SM and DiH SM; C39:0 Triacylglycerol for TAGs. **(B)** Comparison of the relative total lipids in HT-29 cells across control, 4 μM ARC39, necroptosis, and necroptosis + ARC39 conditions. The relative total lipids were quantified by adding the internal standard-corrected abundance for each lipid species per condition which were then normalized to the total lipids of the control. Data represents the mean ± standard deviation; *n* = 4, ns represents *p* > 0.05, * represents *p* < 0.05, ** represents *p* < 0.01, *** represents *p* < 0.001.

Consistent with our previous findings,^10^ ceramides showed a robust and broad increase during necroptosis (**Figure 3A-B**). Notably, ARC39 treatment strongly reduced the necroptosis-associated increase in ceramide levels, resulting to an overall two-fold decrease in total ceramides. In contrast, the levels of sphingomyelins and glycosylated ceramides showed comparatively modest changes. These results indicate that ARC39 exerts its protective effects primarily through suppressing ASMase-dependent ceramide accumulation rather than broadly altering sphingolipid metabolism.

We also examined whether ARC39 treatment affected other major lipid classes (**Figure 3**). Phosphatidylcholines, an abundant membrane phospholipid, showed no significant changes during necroptosis and were not altered by ARC39 treatment, indicating that ASMase inhibition does not broadly disrupt phospholipid composition (**Figure 3A-B**). In contrast, triacylglycerides accumulated during necroptosis, as observed previously, and this accumulation was reduced upon ARC39 treatment (**Figure 3A-B**). Because triacylglyceride accumulation is associated with oxidative stress,^59^ its reversal likely reflects the improved survival of ARC39-treated cells rather than a direct effect on neutral lipid metabolism, further supporting ceramide depletion as the primary contributor to necroptotic rescue.

These results collectively show that the pharmacological inhibition of ASMase reduces necroptosis-induced ceramide accumulation. This change correlates with the observed increase in cell viability and the restoration of membrane integrity. These findings support a model in which ASMase-derived plasma membrane ceramides contribute to membrane permeabilization during necroptosis, and their reduction through ARC39 treatment stabilizes the membrane and attenuates necroptosis.

### ASMase inhibition does not suppress MLKL activation during necroptosis

Ceramides can influence membrane properties, including curvature, fluidity, and lateral organization, microdomains that interface with classical lipid rafts.^18, 60, 61^ These platforms can function as hubs for protein clustering, recruitment, and assembly of signaling complexes. In many forms of regulated cell death, including apoptosis and necroptosis, ceramide-rich membrane domains play roles in amplifying signaling output, reorganizing membrane topology, or facilitating the insertion or stabilization of pore-forming proteins.^18, 62–66^ In necroptosis, ceramide accumulation may influence the biophysical state of the plasma membrane or regulate protein localization in ways that modulate MLKL function or other terminal membrane-disruptive events.

To determine whether the protective effect of ASMase inhibition arises from altered necroptotic signaling upstream of membrane permeabilization, we examined the impact of ARC39 treatment on MLKL expression and phosphorylation. Necroptosis was induced in HT29 cells in the presence or absence of ARC39, and MLKL activation was assessed by collecting whole cell-lysates then immunoblotting for total MLKL and phosphorylated MLKL (pMLKL). Necroptosis induction resulted in robust phosphorylation of MLKL, in agreement with activation of the canonical RIPK1/3-MLKL signaling cascade (**Figures 4A-C, Figure S1-3**).

**Figure 4.**
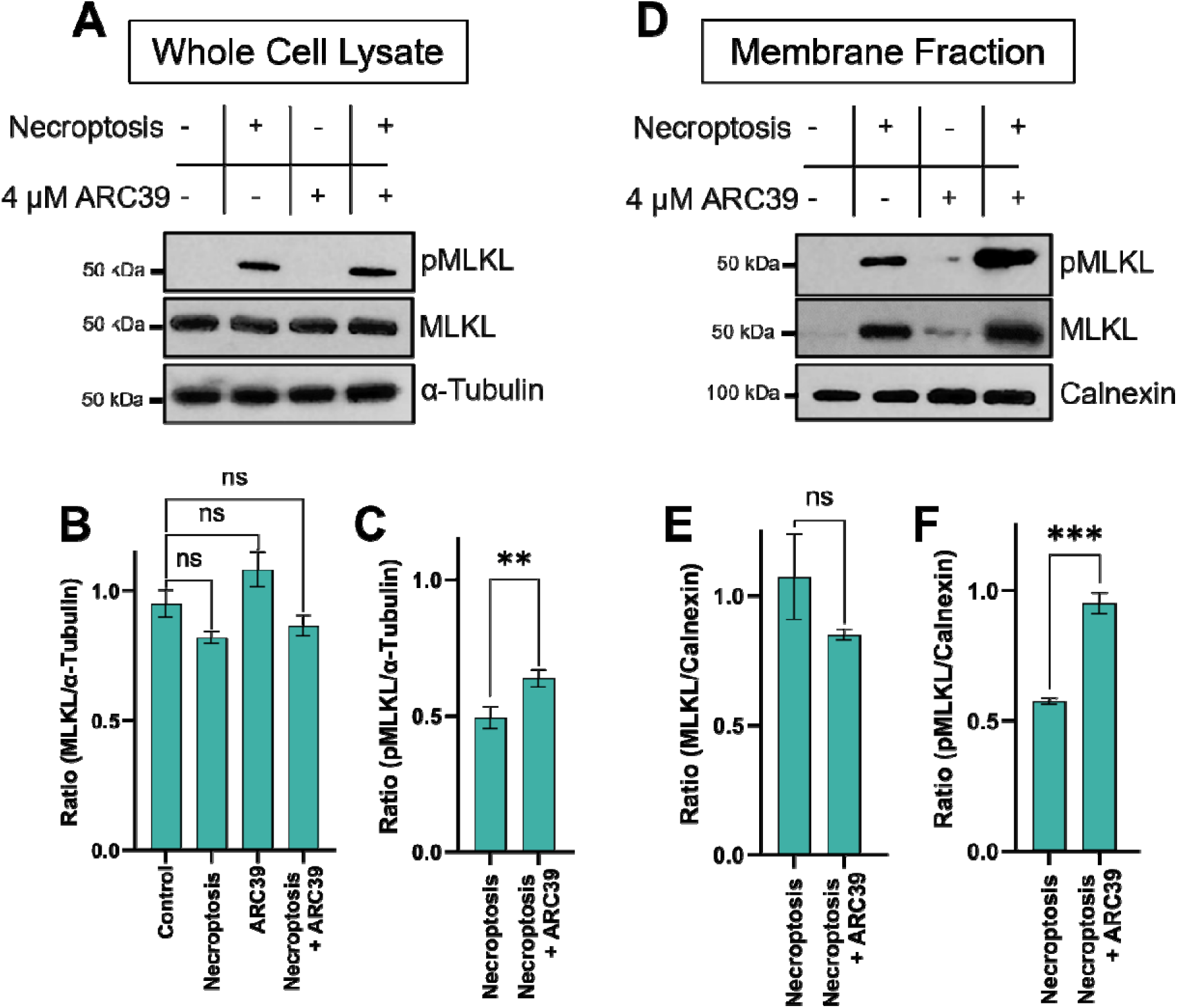
ARC39 does not reduce MLKL phosphorylation during necroptosis. **(A)** Representative Western blot images of whole-cell lysates from HT-29 cells under control, necroptosis, 4 μM ARC39-treated, and necroptotic cells treated with 4 μM ARC39. α-Tubulin was used as the loading control. The raw images used are shown in **Figure S1**. **(B)** Quantification of MLKL levels relative to α-Tubulin in HT-29 cells. The raw images used are shown in **Figure S2**. **(C)** Quantification of pMLKL levels relative to α-Tubulin in HT-29 cells. The raw images used are shown in **Figure S2**. **(D)** Representative Western blot analysis of the membrane fractions of HT-29 cells under control, necroptosis, 4 μM ARC39, and necroptosis with 4 μM ARC39 treatment. Calnexin was used as the loading control. The membrane fractions were produced by ultracentrifugation. The raw images used are shown in **Figure S3**. **(E)** Quantification of MLKL levels relative to Calnexin in HT-29 cells. The raw images used are shown in **Figure S4**. **(F)** Quantification of pMLKL levels relative to Calnexin in HT-29 cells. The raw images used are shown in **Figure S4**. For quantification, the relative intensities of MLKL and pMLKL were calculated using ImageJ by dividing the background-corrected pixel density of the target protein by the background-corrected pixel density of α-Tubulin (whole lysate) or Calnexin (membrane fraction). Data represents the mean ± standard deviation; n = 3, ns represents *p* > 0.05, ** represents *p* < 0.01, *** represents *p* < 0.001.

Notably, ARC39 treatment did not suppress MLKL phosphorylation (**Figure 4A-C**). Instead, pMLKL levels were maintained or modestly increased relative to necroptotic controls. Total MLKL protein levels likewise exhibited a small but significant increase in ARC39-treated cells, indicating that ASMase inhibition does not attenuate MLKL activation and that upstream necroptotic signaling remains intact. To further determine whether ARC39 affects the membrane targeting of activated MLKL, we examined membrane-associated pMLKL levels following subcellular fractionation (**Figures 4D-F**). Consistent with the whole-lysate data, ARC39 treatment did not reduce membrane-associated pMLKL. In fact, phosphorylated MLKL levels within membrane fractions were slightly elevated relative to necroptotic controls. These findings indicate that ARC39 does not affect MLKL phosphorylation or impair the initial translocation of activated MLKL to cellular membranes.

Although MLKL is phosphorylated and activated, its capacity to productively disrupt the plasma membrane appears to be compromised under conditions of ASMase inhibition, based on restoration of membrane permeability (**Figure 2**). Following phosphorylation, MLKL oligomers are recruited to the plasma membrane through interactions with phosphatidylinositol phosphates (PIPs), which serve as interaction platforms that enable initial membrane association.^67^ Disruption of MLKL-PIP interactions impairs membrane targeting and necroptotic execution despite intact upstream signaling, underscoring the importance of lipid cofactors in licensing MLKL function.^68, 69^ In this context, ASMase inhibition may result in defective MLKL-membrane association. While PIPs mediate electrostatic recruitment, additional lipid components can contribute to stabilizing MLKL oligomers and enabling productive insertion. Ceramides are plausible contributors to this step. By promoting membrane condensation and ordered microdomain formation, ceramides may facilitate MLKL insertion, retention, or pore assembly following PIP-mediated recruitment. Thus, under ARC39 treatment, phosphorylated MLKL oligomers may still localize to membranes but fail to execute permeabilization.

### Genetic inactivation of ASMase partially phenocopies ARC39-mediated necroptotic rescue

To validate that the protective phenotype observed with ARC39 treatment arises from inhibition of ASMase, we genetically knocked down ASMase using RNAi. HT29 cells were transduced with shRNA targeting *SMPD1* (shSMPD1), the gene encoding ASMase, and knockdown efficiency was assessed by quantitative ddPCR (**Figure 5A**). This approach resulted in an ∼80% reduction in *SMPD1* transcripts compared to cells expressing a non-targeting control shRNA (shRFP), confirming effective genetic suppression of ASMase expression.

**Figure 5.**
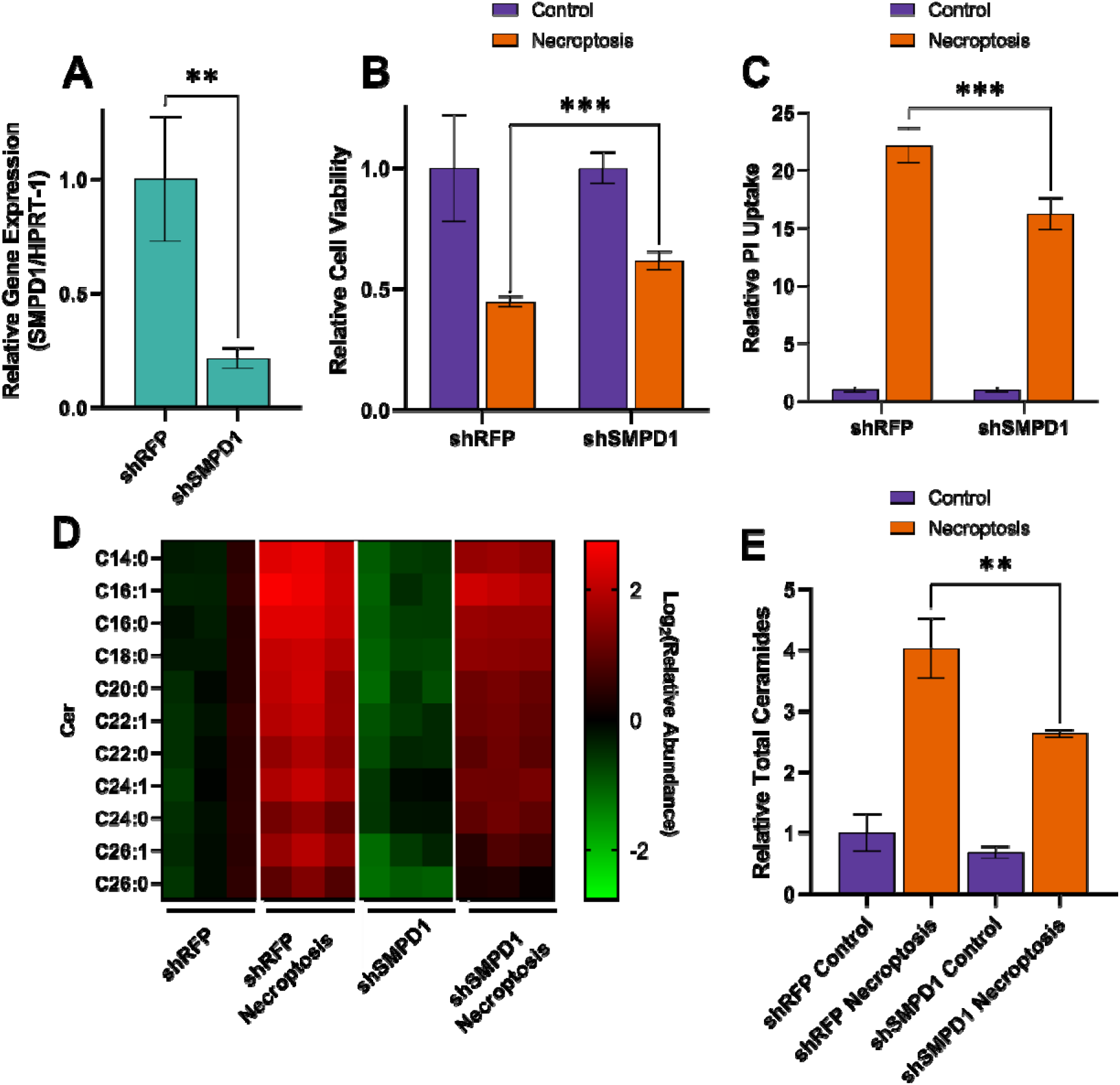
SMPD1 knockdown moderately reduces ceramide levels and partially rescues cells from necroptotic cell death. **(A)** shSMPD1 knockdown efficiency in HT-29 cells. Short hairpin Red Fluorescent Protein (shRFP) was utilized as a negative control. Relative gene expression was calculated by normalizing SMPD1 expression to the housekeeping gene HPRT1. Data represents the mean ± standard deviation; *n* = 3, ** represents p < 0.01. **(B)** Genetic knockdown of SMPD1 increased cell viability compared to the shRFP control under necroptotic conditions in the MTT Assay. Data represents the mean ± standard deviation; *n* = 5, *** represents p < 0.001. **(C)** PI uptake decreased following SMPD1 knockdown, suggesting reduced plasma membrane rupture. Data represents the mean ± standard deviation; *n* = 5, *** represents *p* < 0.001. **(D)** Heatmap showing the relative abundance of ceramides in control vs. necroptosis conditions for shRFP and shSMPD1. Relative abundance was calculated by normalizing each ceramide species to the C17 Cer internal standard. and then to its respective control. Data is shown as Log_2_(relative abundance). Internal standard used for correction: C17:0 Ceramide for Cer **(E)**. Total ceramide levels are reduced upon SMPD1 knockdown. Data represents the mean ± standard deviation; *n* = 3, ** represents *p* < 0.01

We then evaluated the impact of ASMase knockdown on necroptotic cell death. *SMPD1* knockdown resulted in modest restoration in cell viability during necroptosis (**Figure 5B**). More importantly, PI uptake was also reduced in shSMPD1 cells undergoing necroptosis which is consistent with restored plasma membrane integrity (**Figure 5C**). To link the rescue in cell viability and plasma membrane permeability in shSMPD1 cells in necroptosis, we next analyzed the ceramide levels in these cells. ASMase knockdown resulted in a modest decrease of necroptosis-associated ceramide accumulation (**Figures 5D-E**). These findings suggest that the extent of ceramide depletion correlates closely with the degree of necroptotic rescue.

We noted, however, that while knockdown cells exhibited a notable increase in survival relative to control cells, the magnitude of rescue was lower compared to pharmacological inhibition. We propose that the weaker phenotype observed upon genetic ASMase inactivation can reflect adaptive and compensatory remodeling of sphingolipid metabolism that occurs during prolonged genetic perturbation. Because shRNA-mediated knockdown requires extended selection and cellular adaptation period, compensatory pathways may partially restore ceramide pools that are acutely suppressed by ARC39.

Collectively, the genetic data orthogonally validate the pharmacological phenotype observed with ARC39. The convergence of reduced ceramide accumulation decreased membrane permeability and improved survival in both ARC39-treated and shSMPD1 cells, strongly supporting a model in which the necroptotic rescue conferred by ARC39 arises from inhibition of ASMase and consequent changes in sphingolipid profiles, rather than off-target small molecule effects.

## CONCLUSION

Necroptotic execution requires not only activation of the canonical RIPK1/3-MLKL signaling axis but also a membrane environment that permits MLKL to engage and disrupt the plasma membrane. In this study, we identified ASMase-dependent ceramide remodeling as a lipid determinant of this terminal execution step. Pharmacological inhibition of ASMase with ARC39 suppressed ceramide accumulation during necroptosis, restored plasma membrane integrity and cell viability, despite intact upstream signaling and MLKL phosphorylation. Genetic inactivation of *SMPD1* produced a similar albeit weaker phenotype thereby strengthening our proposed model that terminal ASMase-dependent ceramide generation is an important step in necroptotic cell death.

In addition to its role as a lysosomal hydrolase, ASMase can also contribute to plasma membrane remodeling. Upon cellular stress, lysosomal release of ASMase to the plasma membrane occurs, where it hydrolyzes sphingomyelin to generate ceramide-enriched regions. These domains can alter membrane curvature, lipid packing, and organization, and have been implicated in membrane invagination, endocytic clearance of damaged membrane regions, and plasma membrane repair.^70^

In necroptosis, plasma membrane damage is counterbalanced by active repair mechanisms such as ESCRT-III–mediated membrane shedding.^71, 72^ Our findings suggest that ASMase-dependent ceramide generation plays a distinct role in this context. Rather than facilitating membrane repair, necroptosis-associated ceramide accumulation does not appear to promote membrane repair, as membrane-associated pMLKL levels remain unchanged. Instead, ceramide enrichment likely promotes a membrane environment that facilitates efficient MLKL-mediated permeabilization. Specifically, our data support a model in which ASMase-derived ceramide enrichment at the plasma membrane generates a biophysical state that mediates productive MLKL insertion and pore formation. Ceramides promote membrane condensation, alter lipid packing density, and facilitate the formation of ordered microdomains with distinct structural properties.^69^ Such domains may serve as platforms for MLKL oligomer stabilization or conformational changes required for membrane disruption. When ASMase is inhibited, MLKL remains membrane-associated but appears functionally restrained at the level of membrane execution. Thus, necroptosis emerges as a coordinated protein-lipid process in which signaling primes MLKL activation, but membrane lipid remodeling also contributes to the process.

By positioning ceramide remodeling downstream of MLKL phosphorylation, our study reinforces the role of sphingolipids in necroptotic membrane rupture in addition to pore formation by MLKL oligomers. This framework introduces ASMase-dependent sphingomyelin hydrolysis to ceramides as a lipid checkpoint controlling necroptotic sensitivity. Because necroptosis contributes to inflammatory disorders, ischemic injury, neurodegeneration, and cancers, modulation of membrane lipid composition may represent a strategy to tune necroptotic membrane damage without globally suppressing upstream immune signaling pathways.^73^ More broadly, this work highlights membrane ceramide remodeling as an active regulatory layer in necroptosis. However, it is important to note that although the approaches we use to deactivate ASMase reduce ceramide levels and membrane permeability, the precise spatial localization of the functionally relevant ceramide pools was not directly resolved. Future research focus on spatial elucidation of lipid regulation and structural mechanisms linking ceramide remodeling to MLKL-mediated permeabilization remain to be defined.

## SUPPLEMENTAL INFORMATION

Supplemental information includes **Table S1** and **Figures S1-4**. Raw lipid abundances are listed in the *Supplemental Table_lipidomics data* file.

## ACKNOWLEDGMENTS

We acknowledge the support from the NIGMS (R35GM156195 to G.E.A.G.). The authors acknowledge the use of artificial intelligence (AI)-assisted language tools for the purpose of improving grammar, clarity, and overall flow of the manuscript. The AI tools were used solely for language editing. All scientific content, interpretations, and conclusions were developed by the authors.

## AUTHOR CONTRIBUTIONS

Experiments were designed and conducted by L.P., S.C., and G.E.A-G. The study was directed by G.E.A-G. The authors declare no conflict of interest.

## EXPERIMENTAL SECTION MATERIALS

Human colorectal adenocarcinoma epithelial HT-29 cells (Cat # HTB-38) was purchased from the American Type Culture Collection (ATCC). Dulbecco’s Modified Eagle’s Medium (DMEM) (Cat # 10-017-CV), Trypsin (25-050-CI), and Penicillin/Streptomycin (Cat # 30-002-CI) were obtained from Corning. Fetal Bovine Serum (FBS) (Cat # F0926), sodium dodecyl sulfate (SDS) (Cat # L3771), puromycin (Cat # P8833), Polybrene (Cat # TR-1003), ammonium Hydroxide (Cat # 338818), N,N,N′,N′-Tetramethylethylenediamine (TEMED) (Cat # T9281) was purchased from Sigma Aldrich.

LC-MS grade methanol (Cat # MX0486-1), HPLC-grade isopropanol (Cat # PX1834-1) and LC-MS grade acetonitrile (Cat # AX0156-1) was purchased from Millipore Sigma.

Ammonium persufate (APS) (Cat # 0486) was obtained from Thermo Fisher. Chloroform (Cat # 049-1L) was purchased from Honeywell.

Carrier free TNF-α was obtained from R&D systems (Cat # 210-TA/CF). BV6 (Cat # 28830), ZVAD(OH)-FMK (Cat # 14467), FB1 (Cat # 62580), PPMP (Cat # 22677), NVP-231 (Cat # 13858), HPA12 (Cat # 28350), D609 (Cat # 13307), D-NMAPPD (Cat # 10006305), GW4869 (Cat # 13127), DPTIP (Cat # 41681), and LDH Cytotoxicity Assay Kit (Cat # 601170) were obtained from Cayman Chemicals. ARC39 (Cat # 860480) and GT-11 (Cat # 857395) was obtained from Avanti Research. The CellTiter-Glo Luminescent Cell Viability Assay Kit (Cat # G7570) was purchased from the Promega Corporation.

The Micro Bicinchoninic Acid (BCA) Protein Assay kit (Cat # 786-572) was purchased from G-Biosciences.

EDTA-free Protease Inhibitor mini tablets (Cat # PIA32955), Phosphatase Inhibitor Cocktail II (Cat # AAJ61022AA), Pierce™ Bradford Protein Assay Kit (Cat # 23200) and M-PER™ Mammalian Protein Extraction Reagent (Cat # 78501), SuperSignal™ West Pico PLUS Chemiluminescent Substrate (Cat # 34577) and Acrylamide/Bisacrylamide 37.5:1 (Cat # J61505.AP), Propidium Iodide (Cat # P1304MP), MTT reagent (Cat # L11939), Formic Acid (Cat # 270480010), Ammonium Formate (Cat # 168610051) were obtained from ThermoFisher Scientific.

iScript cDNA Synthesis kit (Cat # 1708890), ddPCR Supermix for Probes (No dUTP) (Cat # 1863024), Precision Plus Protein Dual Color Standards (Cat # 1610374), Trans-Blot Turbo 5x Transfer Buffer (Cat # 10026938), 0.2 µm Nitrocellulose membrane (Cat # 1620112), Trans-Blot Turbo RTA Mini 0.2 µm PVDF Transfer Kit (Cat # 1704272) were obtained from Bio-Rad

E.Z.N.A.® HP Total RNA Kit (Cat # D6942) was obtained from Omega Bio-tek, Inc. ESI-L LCMS Tuning Solution (Cat # G1969-85000) was purchased from Agilent.

Antibodies were obtained from Cell Signaling Technology (rabbit monoclonal anti-MLKL, cat. no. 14993S; rabbit monoclonal anti-pMLKL, Cat. #91689), Millipore Sigma (mouse monoclonal anti- α-tubulin, Cat. #T9026) and Jackson Immunoresearch Lab (goat anti-mouse HRP conjugate, Cat # 111-035-144; goat anti-mouse HRP conjugate, Cat # 115-035-174). Autoradiography Film (Cat # CLS-1906-02) was obtained from Chemglass.

C17:0 Ceramide (Cat # 860517P), d^9^ Oleic Acid (Cat # 861809), C17:0 Glucosylceramide (Cat # 860569), C17 Sphingomyelin (Cat # 860656P), and C39 TAGs lipid standards were obtained from Avanti Polar Lipids.

## METHODS

### Cell culture and routine maintenance

HT-29 cells were cultured in DMEM medium that is supplemented with 10% (v/v) FBS and a 1% (v/v) penicillin/streptomycin solution. Cells were grown in a humidified 5% CO_2_ incubator at 37 °C and were routinely monitored under a light microscope for changes in morphology and confluency.

### Necroptosis induction

To induce necroptosis, HT-29 cells were initially sensitized towards TNF-dependent cell death pathways by SMAC mimetic, BV6 (1 μM) and was simultaneously co-treated with pan-caspase inhibitor, zVAD-FMK (25 μM). The cells were then incubated for 30 minutes at 37 °C. After the incubation period, TNF-α (10 ng/mL) was added and the cells were then incubated at 37 °C for 3 hours.

### Small-molecule inhibitor library screen (sphingolipid pathway perturbation)

5 x 10^4^ HT-29 cells were seeded into 96-well plates. After overnight incubation, the inhibitors (originally dissolved in 100% DMSO) were added until the desired concentration was achieved and organic solvent concentration was 0.25%. The control cells received DMSO as vehicle control. After, 21 hours of pretreatment necroptosis was induced as described previously and the final organic solvent concentration after necroptosis induction was 0.75%. For ARC39, the compound was dissolved in 100% PBS and was prepared as described in the literature.^44^ The control cells received PBS as the vehicle control and while the final organic solvent concentration for ARC39-treated conditions was 0.5%

### MTT viability assay

HT-29 cells were plated and pre-treated as described previously.^19, 20^ After pre-treatment and the induction of necroptosis, the 96-well plate was centrifuged at 200 rcf at room temperature for 2 minutes. After centrifuging, the media from each well was removed and 200 μL of fresh media containing 0.5 mg/mL of MTT solution was added into each well. The plate was then incubated for 2 hours. After the incubation period, 155 μL of the media was removed then 90 μL of DMSO was added into each well. The cells were then incubated for 10 minutes, and the resulting formazan crystals was resuspended using a multichannel pipette to break up the formazan crystals. The plate was then centrifuged for 2 minutes at 1000 rcf and the absorbance was read a 550 nm using a plate reader (Biotek Synergy HT). The relative cell viability was calculated by normalizing the blank corrected absorbance of each sample to the average blank corrected absorbance of the control conditions.

### LDH assay

After the treatment period and the induction of necroptosis as described earlier, the LDH release assay was performed using the LDH Cytotoxicity Assay Kit from Cayman Chemicals with minor modifications.^20^ Prior to the assay, the LDH rection solution was prepared as specified by the manufacturer. The 96-well plate was centrifuged at 400 rcf for 5 minutes at room temperature. In a new 96-well plate, 100 μL of the cell supernatant and 100 μL of the LDH reaction solution was added. For the blank, 100 μL of media was added. The plate was then incubated with gentle shaking in an incubator shaker (New Brunswick Scientific C25KC) for 30 minutes at 120 rpm and 37 °C. After the 30 minutes of shaking, the plate’s absorbance was read at 490 nm using a plate reader (Biotek Synergy HT). To calculate the relative LDH release, the blank corrected absorbance of each well was normalized to the blank corrected absorbance of the vehicle control.

### PI uptake assay

After the treatment period and after the induction of necrotic cell death as described previously, the 96-well plate was centrifuged at 200 rcf in room temperature for 2 minutes. After centrifuging, the media from each well was removed and a 200 μL 5 μg/mL PI solution (in PBS) was added into each well. The plate was then incubated for 30 minutes at 37 °C. After the incubation period, the plate was then centrifuged for 2 minutes in room temperature at 200 rcf. The fluorescence intensity was measured using a plate reader (CLARIOstar Plus). The excitation wavelength was measured at 535 nm while the emission wavelength was measured at 617 nm. The relative PI uptake was calculated by normalizing the blank corrected fluorescence of each sample with the average of the blank corrected fluorescence of the control conditions.

### ATP cell viability/Cell Titer Glo assay

The ATP reaction mixture was prepared as specified by the manufacturer. After inducing necroptosis, the 96-well plate was incubated at room temperature for 10 minutes. After the incubation period, the 96-well plate was centrifuged at 200 rcf for 2 minutes at room temperature. 100 μL of media from each well was removed while 100 μL of the ATP reaction mixture was added into each well. The plate was then covered with aluminum foil and gently shaken for two minutes using a microplate vortex mixer (VWR). 150 μL of the cell lysate was then transferred to a new sterile white, solid-flat-bottom, 96-well plate (Promega) and was covered with aluminum foil and allowed to incubate at room temperature for 30 minutes. The luminescence intensity was measured using a CLARIOstar Plus plate reader with the emission wavelength read at 580 nm. The relative cell viability was calculated by normalizing the blank corrected luminescence of each sample to the average of the blank corrected luminescence of the control.

### Lipidomics analysis

1 x 10^7^ *shRFP*-transfected, *shSMPD1*-transfected or wild type HT-29 cells were plated onto 10 cm dishes. *shRFP*- and *shSMPD1*-transfected HT-29 cells were incubated at 37 °C overnight while wild type HT-29 cells are pre-treated with ARC39 as described earlier. The sphingolipid extraction follows a previously published protocol.^20, 57^ Following the treatment period or the induction of necroptosis, the 10 cm dishes are placed on ice and the cells were gently scraped then transferred to 15 mL centrifuge tubes. After scraping, the Plates were rinsed with ∼5mL of cold PBS and was subsequently transferred to the same centrifuge tubes. The cell suspension was then centrifuged for 5 minutes at 500 rcf at 4 °C. After centrifuging, the supernatant was discarded and the resulting cell pellets are washed with cold PBS thrice. The cell suspension was centrifuged and the supernatant was discarded. The cell pellets are then stored at -80 °C for further experiments.

The frozen cell pellets are thawed on ice for ∼30 minutes and are then resuspended in 1 mL of cold PBS. A 50 μL aliquot was then removed and set aside for protein normalization. The remaining 950 μL cell suspension was then transferred into a glass Dounce homogenizer. 1 mL of cold methanol and 2 mL of cold chloroform was then added. The cells were homogenized using a glass pestle with 30 strokes. After homogenizing, the cell pellets were transferred into a 2-dram glass vial and the organic and aqueous layers were separated by centrifuging for 10 minutes at 4 °C and 500 rcf. After centrifuging, the organic layer was carefully transferred into a 1-dram vial using a glass pipette. To ensure that an equal amount of cell material was transferred, 1.5 mL volume was transferred into a new clean 1-dram vial. The chloroform extract was then dried under a stream of inert nitrogen gas (Reacti-Vap™ Evaporator, Thermo Fisher) for approximately 5 minutes per sample. Once dried, the samples normalized by reconstituting with ≥ 150 μL chloroform that is spiked with C17:0 ceramide, d9 Oleic Acid, C17:0 glucosylceramide, C17:0 sphingomyelin and C39 triacylglycerol as internal standards.

Protein content was normalized using the BCA assay following the manufacturer’s protocol. An equal amount of lysis buffer (50 μL) consisting of M-PER Mammalian Protein Extraction Reagent was supplemented with protease inhibitor was added and resuspended to the cell pellets. The cell suspension was kept for 45 minutes for cell lysis. Meanwhile a 1:1 M-PER:PBS buffer solution was prepared and a series of standard solutions of BSA was prepared. Once the cells were lysed, the samples were centrifuged for 15 minutes at 16,900 rcf and 4°C. A 1:10 dilution was then made (diluted with m-per/PBS) and 100 μL of the resulting solution was transferred into a 96-well plate. Afterwards, 100 μL of the BCA reaction mixture was added into each well. The plate was then covered with aluminum foil and gently shaken for 1 hour at 100 rpm and 37 °C. Afterwards, the absorbance of the plate was read at 562 nm. For normalization, a calibration curve was generated from the standard BSA solutions. The volume of was normalized to the lowest protein amount measured.

The LC-MS analysis was performed using an Agilent Infinity 1260 HPLC system that is coupled to an Agilent 6530 Jet Stream ESI-QToF-MS spectrometer. The lipid data was collected over an m/z range of 50-1700 m/z range in extended dynamic range. A DualJSI fitted electrospray ionization (ESI) source was used with the fragmentor voltage set to 175 V and the capillary voltage set to 3500 V. The drying gas temperature was set to 3500 °C and a flow rate of 12 L/min.

Chromatographic separation was achieved using reverse phase gradient chromatography. For negative mode data acquisition, a Gemini C18 reversed-phase column (5μm, 4.6 mm x 50 mm, Phenomenex) with a C18 reversed-phase guard cartridge was used. For positive mode data acquisition, a Luna C5 reversed-phase column (5 μm, 4.6 mm x 50 mm, Phenomenex) C5 reversed-phase guard cartridge was utilized. For both negative and positive mode analysis, a binary solvent system was for gradient elution wherein Mobile Phase A is composed of 95:5 water:methanol (v/v) while mobile Phase B is composed of 60:35:5 isopropanol:methanol:water (v/v). For negative mode analysis, both mobile phase A and B were supplemented with 0.1% (w/v) ammonium hydroxide for improved negative mode detection. Similarly, 5 mM ammonium Formate and 0.1% formic acid (v/v) was instead used in Mobile Phase A and B for improved positive mode detection. The LC gradient was initially at 0% mobile phase B for five minutes then was increased to 100 % mobile phase over 60 minutes. Starting at 65 minutes, an isocratic gradient was applied at 100% B for 7 minutes. Equilibration of the column followed with 0% B for 8 minutes. The flow rate was 0.1 mL/min for the first five minutes and was increased to 0.5 mL/min for the remainder of the gradient.

The lipids of interest were analyzed by extracting the m/z for each lipid species using Masshunter Qualitative Analysis (Agilent, Version 10.0). Each peak was assessed for their identity by making sure that the ppm error was ≤ 20 ppm. The retention time for each lipid species was noted, and the raw abundance was extracted using MassHunter Quantitative Analysis for QQQ (Agilent, version 11.0). The relative abundance was obtained by normalizing each lipid species to the raw abundance of the respective internal standard for each condition then normalizing the internal-standard corrected abundance to the internal-standard corrected abundance of the control conditions. C17:0 Ceramide was used for ceramides, dihydroceramides, and hexosylceramides C17:0 sphingomyelin as used for sphingomyelins and dihydro sphingomyelins while C39:0 triacylglycerol was used for triacyclglycerols. A heatmap was then generated using the Log_2_ of the relative abundance.

### Western Blot

For Western Blot 1 x 10^7^ HT-29 cells are seeded in 10 cm dishes and the cell pellets are collected as previously described. The cell pellets were lysed using with 200 μL of M-PER supplemented with protease inhibitor for 45 minutes on ice. After the incubation period, the cell lysate was centrifuged for 15 minutes at 16,900 rcf, 4°C. The total protein content was determined through the Bradford assay. In a polystyrene cuvette, a 5 μL aliquot of the lysate or lysis buffer solution for the blank, 15 μL of PBS, and 1 mL of the Bradford solution was added. The entire solution was vortexed for ∼5 seconds to ensure homogeneity and was incubated for 7 minutes at room temperature in the dark. The protein concentration was then measured using a Nanodrop 2000 spectrophotometer (Thermo Fisher) at an absorbance of 595 nm.

The samples were separated using a 10% SDS-PAGE gel at 150 V. The separated proteins were then transferred to a nitrocellulose membrane using a Transblot Turbo Transfer System (Bio-Rad) for 7 minutes at constant 1.3 A, 25 V following the manufacturer’s protocol. The membrane was then blocked using 10% nonfat dry milk in tris-buffered saline (TBS)-Tween [10 mM Trisbase, 100 mM NaCl, 0.1% Tween 20 (pH 7.5)] for 1 hour at room temperature. The membranes were then washed thrice in TBS-Tween for five minutes each wash. The corresponding membranes were then incubated at room temperature for 1 hour with their respective primary antibodies (1:2000 dilution for pMLKL and MLKL; 1:10000 for α-tubulin). After incubation, each membrane was quickly washed twice in TBS-tween and was then washed four times with each wash lasting for 5 minutes. The membranes were then incubated at room temperature for 1 hour with secondary antibodies in solution with 5% nonfat dry milk in TBS-Tween. The membranes were then washed using the same way as described earlier and was developed using the using the Super Signal West Pico kit. The western blot image was developed using an autoradiography paper and was briefly washed using a developer and a fixer solution. The resulting x-ray film was scanned in grayscale mode at 600 dpi using a ScanX scanner and was exported as a TIFF, JPEG or PNG image format.

### Western Blot Quantification

Quantification of the Western Blot bands was achieved using ImageJ (version 1.54p; Java 21.0.7 64-bit) using a method as previously described.^20, 74^ Briefly, samples were loaded in triplicate and the western blot was produced as described earlier. ImageJ was opened and the measurement criteria was first defined wherein the set measurements were taken as the mean gray value of the image up to 3 decimal places. The scanned jpeg image was then opened in ImageJ and the pre-selected rectangular tool was used to define the region of interest which is based on the area of the largest band. The resulting selection was saved for each individual analysis. The intensities of each band across a row were measured using the same rectangular frame and the background was measured using the same frame for background corrections. Prior to making background corrections, the pixel density for both sample and background measurements was inverted by subtracting 255 from the measured value. The net protein intensity was then obtained by subtracting the pixel-inverted band intensity of the protein interest from the pixel-inverted band intensity of the background. The relative intensities were then obtained by dividing the corrected pixel density of pMLKL or MLKL with the corrected pixel density of the loading control (α-Tubulin for whole cell lysate; calnexin for membrane fractions).

### Lentiviral Production for SMPD1 Knockdown

Lentiviral particles targeting *SMPD1* were generated using a short hairpin RNA (shRNA) construct cloned into the pLKO.1 backbone (*shSMPD1*: NM_000543, TRCN0000049014), obtained from MilliporeSigma as bacterial glycerol stocks. Bacterial cells were collected from the glycerol stocks using a sterile pipette tip, resuspended in LB medium, and incubated with shaking for 2 h prior to the addition of ampicillin (100 µg/mL). Cultures were grown overnight at 37 °C with agitation. For long-term storage, glycerol stocks were prepared by mixing equal volumes of bacterial culture and sterile-filtered 50% (v/v) glycerol in nanopure water and stored at −80 °C. The remaining cultures were centrifuged at ∼3000 × g for 20 min at 4 °C, and bacterial pellets were stored at −80 °C until plasmid isolation. Plasmid DNA was purified using the E.Z.N.A. FastFilter Plasmid DNA Mini Kit (Omega Bio-tek) following the manufacturer’s protocol.

For lentiviral packaging, approximately 3 × 10 HEK-293T cells were seeded into 6-well plates and allowed to adhere for 16 h. Transfection mixtures were prepared in 150 µL of serum-free DMEM containing 100 ng VSV-G envelope plasmid, 900 ng psPAX2 packaging plasmid, and 1 µg of the transfer plasmid (*shRFP* or *shSMPD1*). X-treme GENE 9 transfection reagent (6 µL) was added to the DNA mixture, which was gently mixed and incubated at room temperature for 20 min. Shortly before transfection, cell culture media were replaced with fresh medium supplemented with 25 µM chloroquine. The transfection complexes were added dropwise to the cells, followed by gentle rocking of the plate to ensure uniform distribution. Cells were maintained for 48 h, after which the viral supernatant was collected, passed through a 0.45 µm syringe filter, aliquoted, and stored at −80 °C.

### Genetic inactivation of A-SMase (SMPD1) in HT29 cells using lentiviral shRNA

For lentiviral transfection, 3 x 10^5^ HT-29 cells were seeded in 6-well plates (n = 5) and were placed inside the incubator for overnight attachment. After overnight attachment, the media was removed and was replaced with 2 mL of fresh media containing 1:2000 polybrene solution. The polybrene solution was allowed to incubate for 5 minutes at room temperature and 100 μL of *shRFP* or *shSMPD1* viral particles were then added into 5 wells. No virus was added into the 6^th^ well to serve as the control. The 6-well plate was then allowed to incubate for two days. After 48-hour incubation, the cells are trypsinized then pooled into a 10 cm dish containing media with 2 μg/mL of puromycin for cell selection. After 2 days, the media was replaced with fresh media containing 1 μg/mL of puromycin and the cells were continuously maintained in media containing 1 μg/mL of puromycin.

### ddPCR

*shSMPD1*- and *shRFP*-transfected HT-29 cells were plated into 10cm dishes for overnight attachment and the cell pellets were collected. The total RNA from these cells were extracted and purified using the E.Z.N.A.® HP Total RNA Kit (Omega Bio-tek) following the manufacturer’s protocol. The total RNA concentration was measured using a Nanodrop-1000 spectrophotometer (Thermo). The extracted RNA was converted into cDNA using the iSCript cDNA synthesis kit and a Biorad Droplet Digital PCR QX200 system. The reaction mixtures for the digital droplet PCR (ddPCR), was prepared in a 96-well plate using the ddPCR Supermix for probes following the manufacturer’s instructions. Water-oil emulsion droplets were generated and transferred into a new 96-well plate using a BioRad automated droplet generator. The new plate was heat-sealed with aluminum foil and was subjected to the PCR reaction. The PCR cycle is as follows: 95 °C for 10 min, followed by 40 cycles of (1) 94 °C for 30 s and (2) 56 °C for 60 s, with a final 10 min inactivation step at 98 °C. After the PCR reaction, a QX200 droplet reader (Biorad) was used to measure and quantify the reference gene (*HPRT1*) the target gene (*SMPD1* or *shRFP*). *SMPD1* primer sequence: sense: 5’-GAGAGAGATGAGGCGGAGA-3’; antisense: 5’- CTGGCTCTATGAAGCGATGG-3’. *HPRT1* primer sequence: sense: 5’-GTATTCATTATAGTCAAGGGCATATCC-3’; antisense: 5’- AGATGGTCAAGGTCGCAAG-3’. Data analysis was done using QuantaSoft software (Biorad).

