## Supplemental data file for "Inhibition of Acid Sphingomyelinase Links Sphingolipid Remodeling to Necroptotic Cell Death"

**Table S1**. Structure of the various Sphingolipid biosynthesis inhibitors that were used in our study. Their corresponding enzyme targets are listed. Inhibitors highlighted in green, orange, and blue represent inhibitors that target enzymes involved in de novo biosynthesis, downstream Sphingolipid, and Sphingolipid breakdown, respectively.

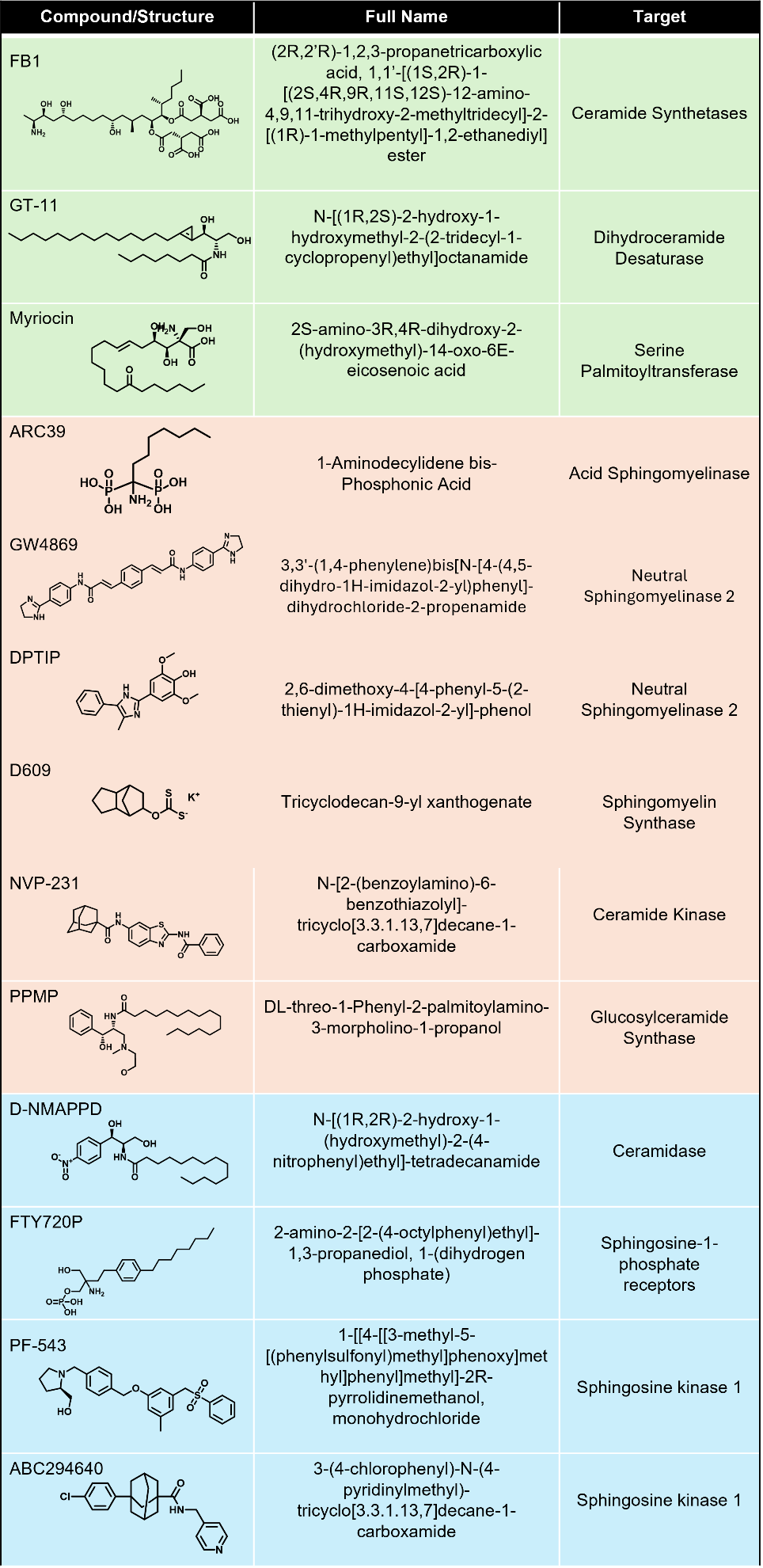

**Table S2. (separated file)**

Targeted lipidomics results. Ion counts for each lipid are provided. Relative abundance was calculated as the ratio of each lipid’s abundance to the mean abundance of the control group at the corresponding time point. This is provided as a separate excel sheet.

Excel File: Table S2

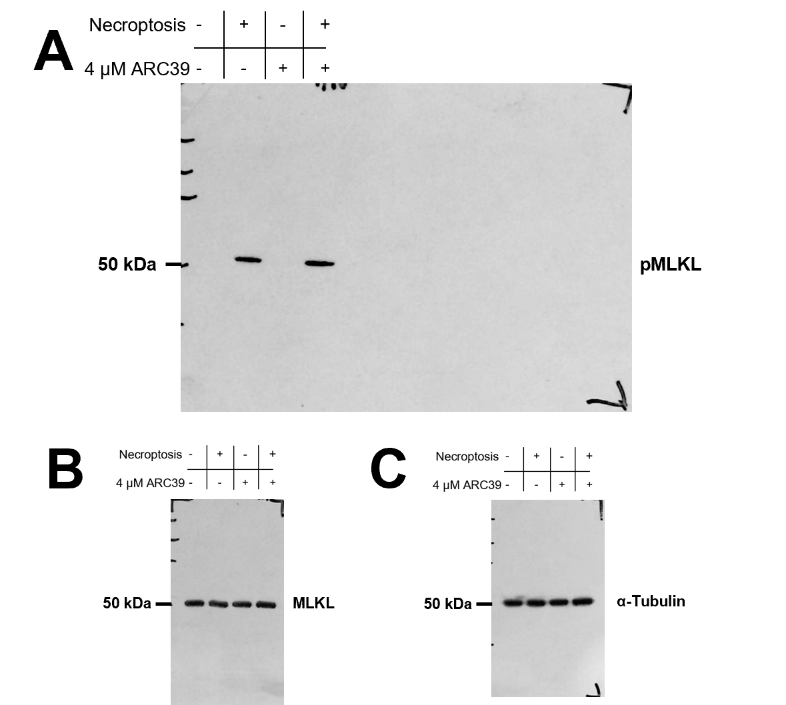

**Figure S1**. Raw images used for the representative Western Blot of the whole cell lysate in Figure 4A. **(A)** Raw image used for pMLKL. **(B)** Raw image used for MLKL. **(C)** Raw image used for α-Tubulin.

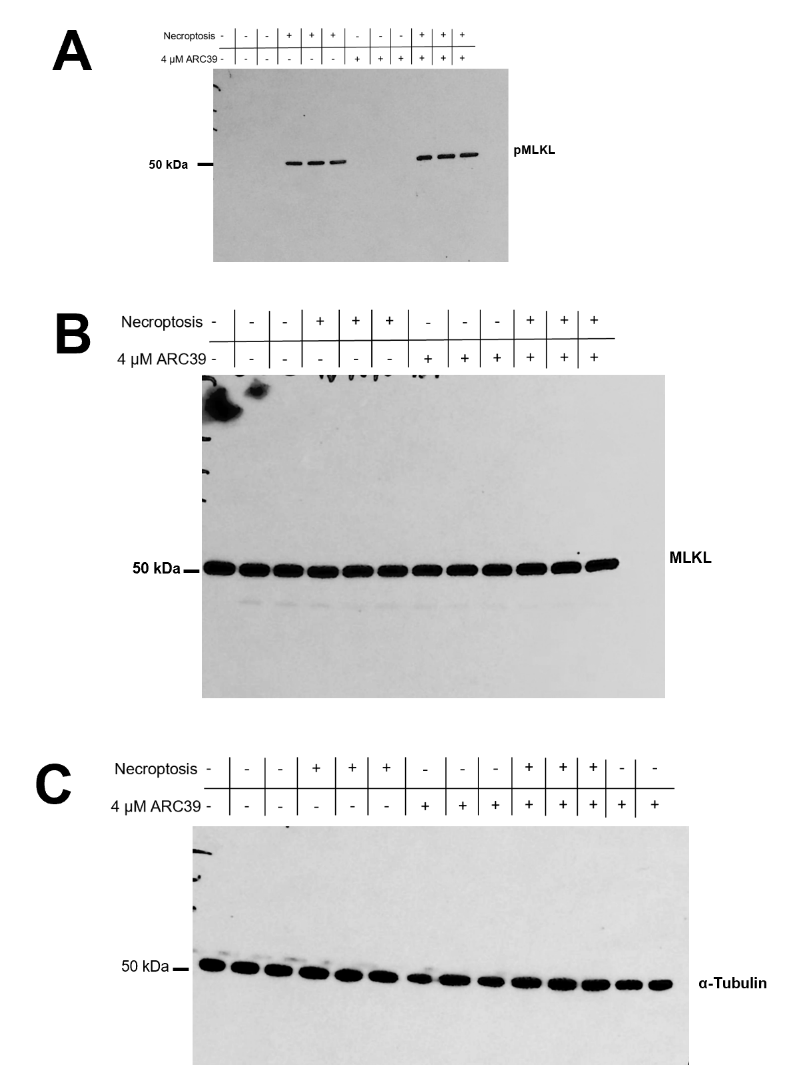

**Figure S2.** Raw images used for the western blot quantification of the whole cell lysate in Figures 4B-C. **(A)** Raw image used for the quantification of pMLKL. **(B)** Raw image used for the quantification of MLKL. **(C)** Raw image used for the quantification of α-Tubulin. Lanes 8, 14 and 15 were used to measure the band intensity of the Necroptosis |-| / ARC39 |+| condition for α-Tubulin.

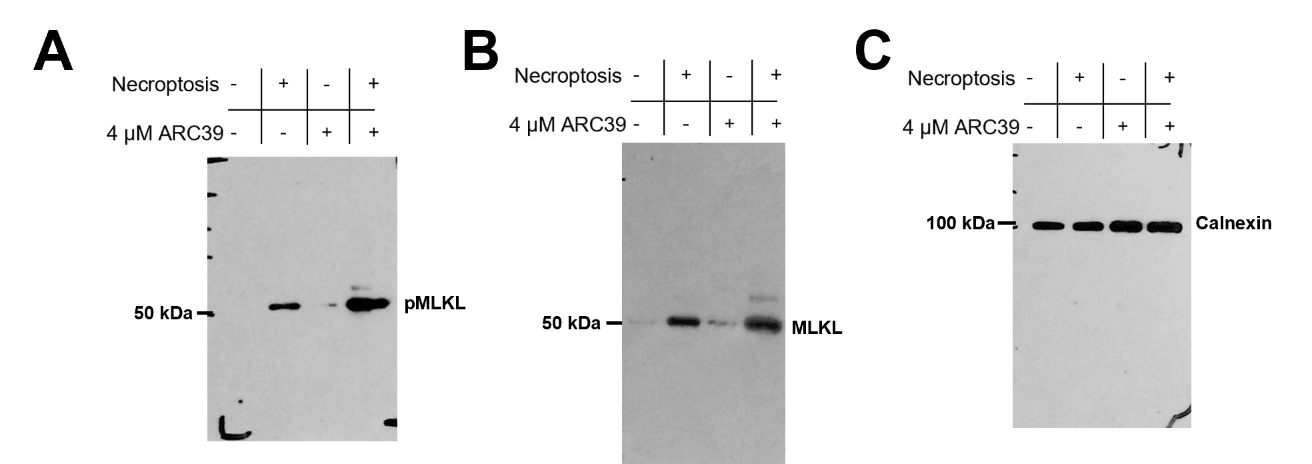

**Figure S3**. Raw images used for the representative Western Blot of the membrane fraction in Figure 4D. **(A)** Raw image used for pMLKL. **(B)** Raw image used for MLKL. **(C)** Raw image used for Calnexin.

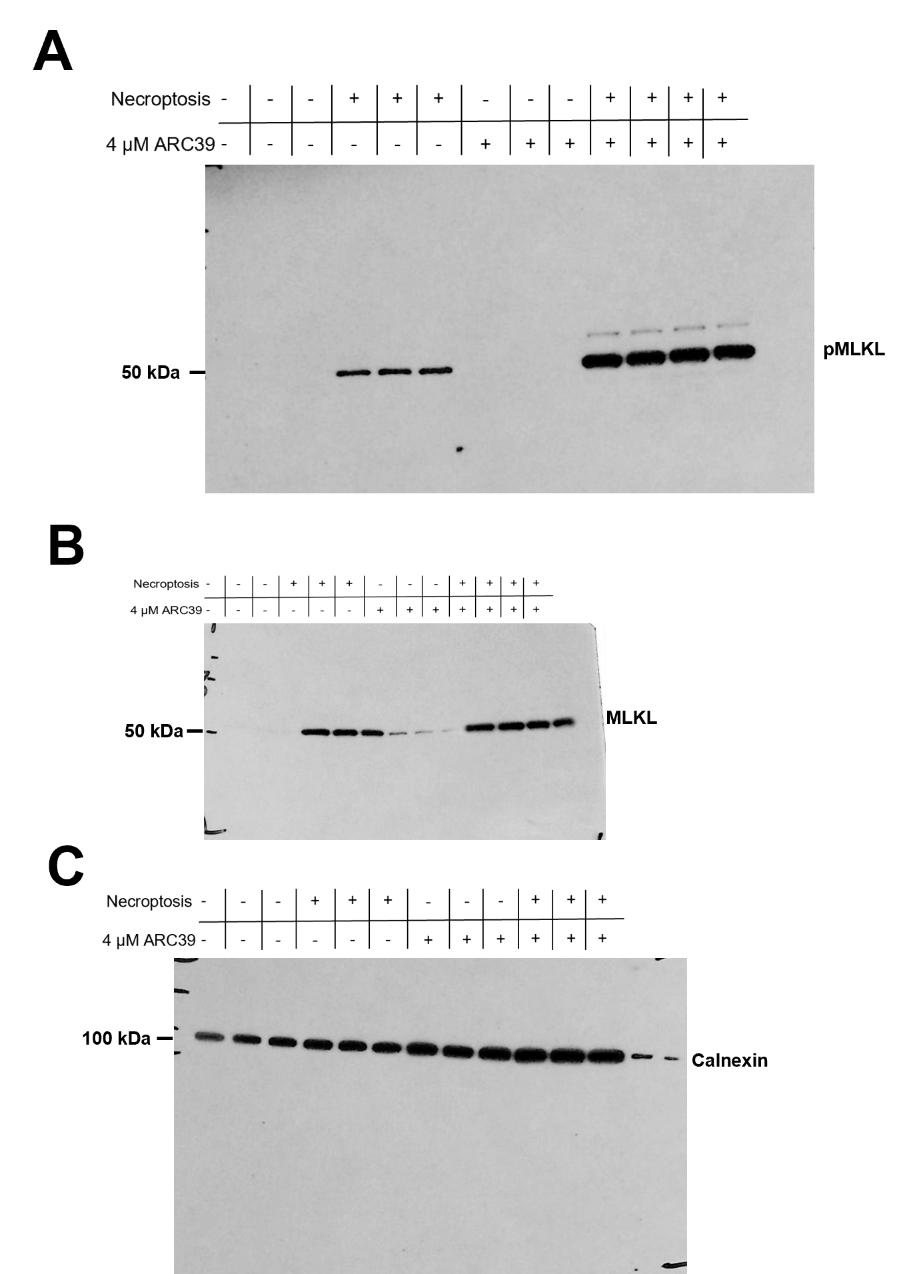

**Figure S4.** Raw images used for the western blot quantification of the membrane fractions in Figures 4E-F. Lanes 10-12 were used for the quantification of the Necroptosis |+| / ARC39 |+| condition for pMLKL, MLKL and Calnexin. **(A)** Raw image used for the quantification of pMLKL. **(B)** Raw image used for the quantification of MLKL. **(C)** Raw image used for the quantification of Calnexin.
